# Vascular Destabilization and Pericyte Detachment are Mediated by hIAPP Aggregation in Transgenic Mice

**DOI:** 10.64898/2026.02.21.707196

**Authors:** J Koepke, L Mateus Gonçalves, C Andrade Barboza, AC Aplin, DJ Hackney, SA Gharib, O Mohn, M Teng, JJ Castillo, J Almaça, RL Hull-Meichle

**Affiliations:** Department of Cell Biology and Alberta Diabetes Institute, University of Alberta, Edmonton, Canada; Division of Endocrinology, Diabetes and Metabolism, Department of Medicine, University of Miami Miller School of Medicine, Miami, FL; Diabetes Research Institute, University of Miami Miller School of Medicine; Department of Veterans Affairs Puget Sound Health Care System, Seattle, WA, USA; Seattle Institute for Biomedical and Clinical Research, Seattle, WA, USA; Division of Pulmonary, Critical Care, and Sleep, Department of Medicine, University of Washington, Seattle, WA, USA; Division of Metabolism, Endocrinology and Nutrition, Department of Medicine, University of Washington, Seattle, WA, USA

**Keywords:** Islet, pancreas, amyloid, IAPP, pericyte, endothelial cell, capillary, type-2 diabetes

## Abstract

**Aims/hypothesis:** Human islet amyloid polypeptide (hIAPP) deposition is a common feature of type-2 diabetes (T2D). Previous studies have demonstrated hIAPP-mediated endothelial cell (EC) dysfunction and inflammation, but little is known about islet microvascular stability or pericyte function in hIAPP-containing islets. This study investigates how islet endothelial cells and pericytes are influenced by hIAPP aggregation.

**Methods:** Bulk RNAseq and qPCR were conducted on hIAPP or vehicle treated MS-1 cells and bead-purified human islet CD31+ cells from donors with or without T2D to determine how islet ECs respond to hIAPP exposure. Confocal imaging of living pancreatic slices obtained from hIAPP transgenic mice was conducted to evaluate the effect of hIAPP deposition on islet pericyte function and vasomotor responses.

**Results:** hIAPP-treated MS-1 cells and ECs purified from T2D islets demonstrate downregulation of leading-edge genes associated with extracellular matrix and cell adhesion pathways. Pericytes from hIAPP-expressing mouse islets appear detached from underlying endothelial cells, which was associated with impaired vasomotor responses to constrictive or dilatory stimuli.

**Conclusions/interpretation:** hIAPP induces vascular destabilization by downregulating mRNA of key extracellular matrix and cell adhesion molecules in ECs, likely promoting the breakdown of EC-EC and EC-pericyte coupling. hIAPP disrupts EC-pericyte connections, and pericyte detachment ultimately impairs pericytes’ ability to modulate capillary diameter without impairing intracellular Ca^2+^ dynamics. Our data suggest that amyloid deposition compromises EC health and survival by altering islet microvascular morphology, stability, and function. This, in turn, may disrupt islet microvascular stability and exacerbate endocrine cell dysfunction in T2D.

**Research in context:** *What is already known about this subject?:* - hIAPP is cytotoxic to islet endothelial cells and beta cells, and contributes to islet failure in type-2 diabetes (T2D) - hIAPP transgenic mice demonstrate islet capillary dilation, loss of vascular structures, and increased pericyte density - Impaired pericyte anchorage and vascular fragmentation drive diabetes-related vasculopathies in other tissues, like the retina, kidney, and brain

*What is the key question?:* - How does the surviving microvasculature in islets respond to hIAPP deposition?

*What are the new findings?:* - Endothelial cells demonstrate transcriptional downregulation of key genes involved in cytoskeleton, ECM, and cell-adhesion maintenance, including *Thbs1*, *Tln1*, and *Plec*. - Amyloid deposits disrupt homeostatic interactions between endothelial cells and pericytes. - Amyloid-adjacent islet pericytes are detached from endothelial cells and display impaired ability to modulate capillary diameter.

*How might this impact on clinical practice in the foreseeable future?:* - Therapies targeting endothelial cell-pericyte interactions may restore islet microvascular stability and improve islet function, especially in the context of early T2D.

## INTRODUCTION

Pancreatic islets are highly vascularized, with capillaries not only perfusing endocrine cells with nutrients and growth factors but also providing extracellular matrix (ECM) and signalling components that are vital for islet homeostasis [1–5]. Islet capillaries are comprised of fenestrated endothelial cells (ECs) and contractile pericytes [6]. Microvascular tone is controlled by pericytes, which constrict or dilate capillaries in response to extracellular signals like plasma glucose, beta cell-derived adenosine, or EC-derived endothelin-1 (ET-1) [5, 7]. Pericytes-EC contact is essential for maintaining vascular integrity and selective permeability in specialized capillary beds, where failure to maintain proper pericyte coverage impairs capillary function in several pathologies [8–10]. Little is known about changes in islet pericyte coverage during the pathogenesis of T2D or its relative contribution to metabolic homeostasis. In mouse models of type-2 diabetes (T2D), islet capillaries become leaky and poorly responsive to vasoactive stimuli, impairing islet perfusion and contributing to a hostile, proinflammatory extracellular environment that exacerbates islet dysfunction [7, 11–14]. In T2D human islets, endocrine cell dysfunction is accompanied by histological evidence of islet microvascular dysfunction [15]. While T2D microvascular dysfunction is well characterized in the retina and kidney of individuals with T2D, islet microvasculopathies remain an emerging area of research interest [16].

Islet Amyloid Polypeptide (IAPP) is an endogenous beta cell peptide that is co-secreted with insulin [17]. Under certain conditions, human IAPP (hIAPP) can form extracellular aggregates that elicit cytotoxic effects on neighbouring cells [18]. There is evidence of hIAPP deposition in a majority of autopsy samples from humans with T2D [19, 20], where amyloid deposits are associated with beta cell loss and vascular remodelling [15, 20–22]. While rodent IAPP is non-amyloidogenic, transgenic expression of hIAPP recapitulates its cytotoxicity in mice. Mouse islets with transgenic hIAPP expression in combination with high glucose exhibit increased beta cell apoptosis [23, 24]. Furthermore, transgenic mice expressing hIAPP can develop islet amyloid deposits and impaired glucose tolerance [25], along with transcriptional changes consistent with those observed in human T2D islets [26]. Beyond islet endocrine impairment, hIAPP TG mice exhibit markers of endothelial inflammation and altered capillary morphology [13].

Previous studies have demonstrated direct hIAPP-induced cytotoxicity in cultured islet ECs, while hIAPP transgenic mice exhibit decreased capillary density, vascular inflammation, and amyloid deposition between beta cells and neighbouring capillaries [13, 25]. Despite these observations, it is unclear how remaining ECs respond to amyloid deposition, or how such changes impact islet microvascular function. In the present study, we investigate how surviving ECs respond to hIAPP exposure and the functional impact of hIAPP deposition on microvascular tone and pericyte contractility.

## METHODS

### Human islet CD31+ cells and MS-1 cells

Human islets were obtained from organ donors with and without T2D through the Integrated Islet Distribution Program (donor information is summarized in **Suppl Table 1**). Islets were dispersed immediately upon receipt, and islet ECs were isolated by anti-CD31 conjugated magnetic bead separation (Miltenyi Biotec, USA) as previously described [13]. Islet preparations with post-isolation culture time <24 hours before shipping were prioritized to maximize survival of intra-islet ECs, which die off rapidly after islet isolation [27]. Enrichment of CD31+ cells was confirmed by qPCR (**Suppl Fig 1**).

MS-1 cells (mouse islet ECs; CVCL_6502) were maintained in complete culture media at 37 °C in a 5% CO_2_ humidified incubator. Media was comprised of RPMI 1640 with 11.1 mmol/l glucose, 10% FBS, 1 mmol/l sodium pyruvate, 100 U/ml penicillin, and 100 μg/ml streptomycin. All MS-1 assays were conducted in complete culture media with added vehicle or drug treatment as described below.

### Preparation of IAPP and treatment of MS-1 cells

Human IAPP peptide was obtained from Dr. Daniel Raleigh, reconstituted in 1-1-1-3 hexafluoroisopropanol and lyophilized in 100 μg aliquots. IAPP was then resuspended in Tris-HCl buffer (20 mmol/l, pH 7.4) and diluted into complete media at a final concentration of 20 μmol/l immediately prior to use.

cDNA from existing MS-1 cell samples was also analyzed in this study. Briefly, MS-1 cells were cultured with hIAPP in the presence or absence of DMSO vehicle (0.1% vol./vol.), the amyloid aggregation inhibitor Congo Red (200 μmol/l), or the TLR2/4 inhibitor OxPapC (30 μg/ml) as described previously [13].

### RNA isolation and sequencing

*Bulk sequencing -* Total RNA was extracted from human CD31+ and MS-1 islet EC samples and sequenced by the Benaroya Research Institute Genomics Core. We conducted DESeq2 analysis to identify differentially expressed genes using Benjamini-Hochberg adjusted p-values [28]. Gene Set Enrichment Analysis (GSEA) using Gene Ontology (GO) annotation and leading-edge gene analysis was performed to identify enriched processes and their main contributors [29]. Significantly enriched gene sets were identified by permutation-based false discovery rate (FDR). Network analysis and visualization of interacting GO gene sets was performed using Cytoscape (v 3.8.0).

*qPCR -* RNA was harvested (High Pure RNA Kit; Roche) and reverse transcribed (High-Capacity cDNA Reverse Transcription Kit; ThermoFisher). cDNAs were analyzed in triplicate, using prevalidated TaqMan primer-probes (**Suppl Table 2**). Results were calculated using PPIB (human samples) or 18S ribosomal RNA (MS-1 samples) as the housekeeping gene, with the 2−ΔΔCt method.

### Mouse model generation, diet, and experimental timeline

Mouse work was approved by Seattle VA Puget Sound Health Care System’s Institutional Animal Care and Use Committee. Live pancreas slice work was approved by the University of Miami Institutional Animal Care and Use Committee (Miami, FL).

Male C57BL/6xDBA/2J background mice, with or without hemizygous transgenic expression of human islet amyloid polypeptide (TG or NT, respectively), were fed a diet containing 18% kcal from fat (PicoLab Mouse Diet 20 5058; Purina Lab Diet) for at least 12 months, to stimulate hIAPP aggregation [30]. Female mice were not included in this study since only ∼10% of female TG mice develop amyloid, compared to ∼80% of male TG mice after a year of high-fat feeding [25].

10 mice (N=7 TG, N=3 NT) underwent oral glucose tolerance testing before transport to the University of Miami. Briefly, mice were fasted for 6 hours, and baseline blood glucose levels were determined by tail bleed. Thereafter, glucose (2 g/kg) was administered by oral gavage, and blood samples were collected at 10, 20, 30, 60, 90, and 120 minutes. Blood glucose was measured using an Accu-Chek Aviva Plus glucometer.

### Lectin infusion and slice generation

150 μm thick living pancreas slices were prepared from TG and NT mice injected with lectin (DyLight 649; 70 μL) 10 min before euthanasia as previously described [7, 31]. Slices were loaded with 6 µmol/L Fluo4 for 60 minutes in 3mmol/L glucose Hepes buffer before Ca^2+^ imaging with an upright confocal microscope (Leica SP8). We performed 3-5-minute time-lapse recordings at 1 frame per second to record changes in islet pericyte [Ca^2+^]_i_ and blood vessel diameter induced by endothelin-1 (10 nmol/L), bosentan (100 nmol/L), angiotensin-II (100 nmol/L), glucose (16 mmol/L), and epinephrine (10 μmol/L). Islet ROIs were identified by tissue backscatter using excitation at 638 nm and emission at 650-750nm, as previously described [31]. At the time of imaging, pericytes were distinguished from smooth muscle cells based on their morphology and location within the islet. Pericyte identity was confirmed through *post hoc* NG-2 staining after slice fixation as described below.

### Image analysis

Living pancreas slices were immersed in 4% paraformaldehyde for 1 hour and washed with PBS. Slices were incubated overnight with NG-2 (1:100; AB5320; EMD Millipore) and CD31 primary antibodies (1:25; #550274; BD Pharmigen). Samples were counterstained with AlexaFluor goat anti-rabbit or anti-rat antibodies, and Thioflavin-S (1% wt/vol.; Sigma-Aldrich). Confocal images of islets were taken every 3–4 μm, spanning ∼40 μm of the islet, and Z-projections were analyzed in ImageJ to quantify the immunostained area for each antigen and normalized by dividing by the islet ROI area. Data are presented as the average value per islet analyzed.

In separate animals, Dylight 649-conjugated Lycopersicon esculentum (2 mg/ml; Vector Laboratories) was infused IV perimortem, followed by 10% neutral buffered formalin to preserve tissue morphology. 30 μm pancreas sections were stained with NG-2 (1:100; AB5320), AlexaFluor 586 goat anti-rabbit (1:200; A11011; Invitrogen) and Thioflavin-S (1% wt/vol). Confocal images were acquired using a 40X oil objective (NA 1.3) with Nyquist sampling on a resonance-scanning head (A1R-HD; Nikon).

### Statistics

Statistical analysis was conducted in GraphPad Prism 10. All data are presented as mean ± SEM, with individual datapoints superimposed. For comparisons between two groups, we used a Welch’s two-tailed t-test. For comparisons between three groups, we used an Ordinary one-way ANOVA with correction for multiple comparisons. Statistical significance is denoted by asterisks where * p<0.05, ** p<0.01, *** p<0.001, and ****p<0.0001.

## RESULTS

To investigate how surviving ECs respond to hIAPP-induced cytotoxicity, we compared the transcriptional profile of immortalized mouse islet endothelial MS-1 cells treated with vehicle or hIAPP for 48 hours *in vitro*. In parallel, we collected human islet CD31+ ECs from organ donors with and without T2D to assess differential gene expression. We found over 1,500 significantly differentially expressed genes in MS-1 cells (adjusted p-value <0.01), but none in human CD31+ cells after correction for multiple testing [**Suppl Fig 2**]. To compare both models at the pathway level, we conducted Gene Set Enrichment Analysis (GSEA) to investigate differentially regulated Gene Ontology (GO) gene sets. MS-1 cells demonstrated both up- and downregulated GO processes in response to hIAPP treatment. Processes associated with gene transcription, DNA repair, and cell cycle regulation were increased with hIAPP treatment, while pathways associated with extracellular matrix remodelling and metabolism were decreased [**Fig 1A**]. Similarly, CD31+ cells from T2D islets exhibited downregulated GO processes, mostly associated with ECM remodelling, cell adhesion, the inflammatory response, metabolism, and cell migration [**Fig 1B**].

**Figure 1:**
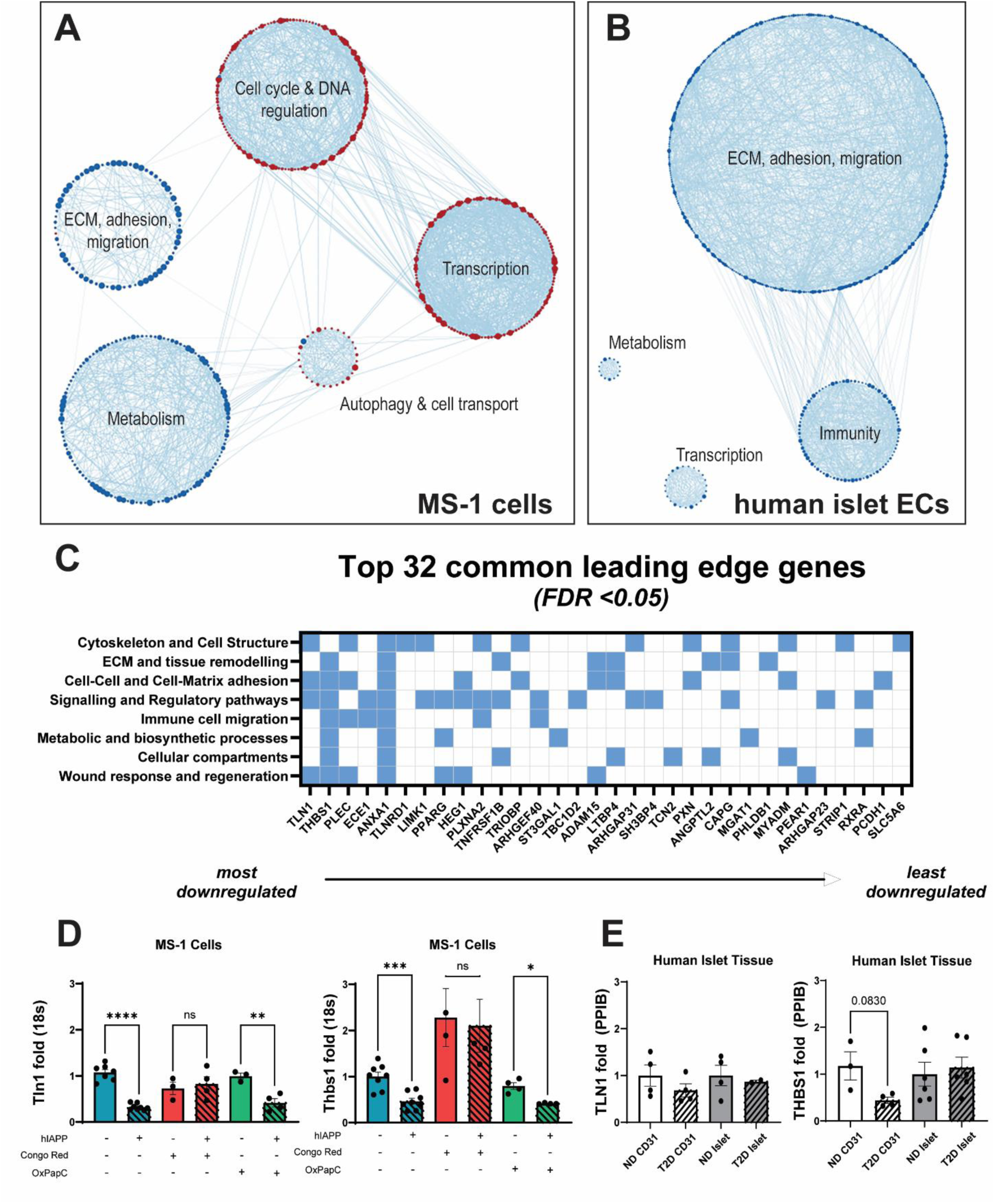
Gene Set Enrichment Analysis and Leading Edge Genes. Network representation of Gene Set Enrichment Analysis (GSEA) from hIAPP-treated vs untreated MS-1 cells (A) or islet CD31+ cells from human samples with or without type-2 diabetes (T2D) (B). Each node represents an upregulated (red) or downregulated (blue) Gene Ontology (GO) geneset (FDR < 0.05). Gene sets sharing 50% or more common genes are connected, resulting in the emergence of large biological modules with distinct functional themes. C) Leading-edge genes common between MS-1 and human CD31+ cell datasets, filtered by endothelial cell-specific GO pathways. D) qPCR of the top two leading edge genes in MS-1 treated with 20µmol/l hIAPP in the presence or absence of the amyloid aggregation inhibitor Congo Red (200 μmol/l) or TLR2/4 inhibitor OxPapC (30 μg/ml). E) qPCR of the top two leading edge genes in bead-purified islet CD31+ cells or unsorted human islets from non-diabetic (ND) or T2D tissue donors. qPCR data were analyzed by Welch’s t-test, where statistical significance is denoted by asterisks (* p<0.05, ** p<0.01, *** p<0.001, and **** p<0.0001).

Next, we conducted leading-edge gene analysis, focusing on enriched GO gene sets and common transcripts in both hIAPP-treated MS-1 cells and T2D human islet CD31+ cells. This comparison allowed us to identify transcriptional changes relevant to human islet ECs in T2D, focusing on hIAPP-dependent effects rather than broader aspects of T2D pathophysiology. Our analysis revealed 35 common downregulated leading-edge genes across 276 human and 98 mouse GO gene sets [**Suppl Table 3**]. We categorized each common or equivalent GO processes into groups representing different aspects of EC function, ultimately identifying 32 common genes deemed relevant to vascular stability. These GO pathway categories included maintenance of EC function (ex. cytoskeleton and cell structure, signalling and regulatory pathways) and maintenance of the extracellular environment (ex. ECM and tissue remodelling, cell-cell and cell-matrix adhesion) [**Fig 1C, Suppl Fig 3**]. From this curated list of 32 leading-edge genes, *Thbs1* and *Tln1* emerged as genes of interest for further investigation in MS-1 cells.

To confirm that these genes were indeed influenced by hIAPP aggregation, we reanalyzed existing MS-1 cell cDNA samples from cells treated with hIAPP in the presence of the amyloid blocker Congo Red or the TLR2/4 blocker OxPaPc. We found that *Thbs1* and *Tln1* were downregulated by hIAPP, with this transcriptional change attenuated by Congo Red but not OxPaPc [**Fig 1D**]. Complementary qPCR analysis was conducted in human islet lysate and bead-purified CD31+ cells, although neither transcript reached statistical significance in human tissue [**Fig 1E**]. Overall, these findings confirm that downregulation of *Thbs1* and *Tln1* mRNA is dependent on hIAPP aggregation rather than TLR-mediated inflammation in MS-1 cells.

Our transcriptional characterization of islet ECs suggested impairment of cell structure, ECM maintenance, and cell adhesion. As such, we investigated how islet EC-pericyte interactions are affected by hIAPP deposition using a transgenic mouse model. Male hIAPP transgenic (TG) mice raised on a high-fat diet for an average of 52 weeks (50-56 weeks) demonstrate expectedly heterogeneous hIAPP aggregation and varying levels of glucose tolerance [**Suppl Fig 4**], consistent with previously published studies in these mice. Furthermore, these mice exhibited decreased islet backscatter, a measure of granularity, and altered Ca^2+^ dynamics [**Suppl Fig 4**], making them a suitable model of islet impairment. High-magnification volume projections of islet vasculature in TG and non-transgenic littermates (NT) revealed striking differences in EC-pericyte associations. Amyloid-adjacent pericytes were dissociated from neighbouring ECs such that hIAPP deposits formed between the two cell types, while NT pericytes were closely associated with neighbouring ECs [**Fig 2, Suppl Movie 1-3**].

**Figure 2:**
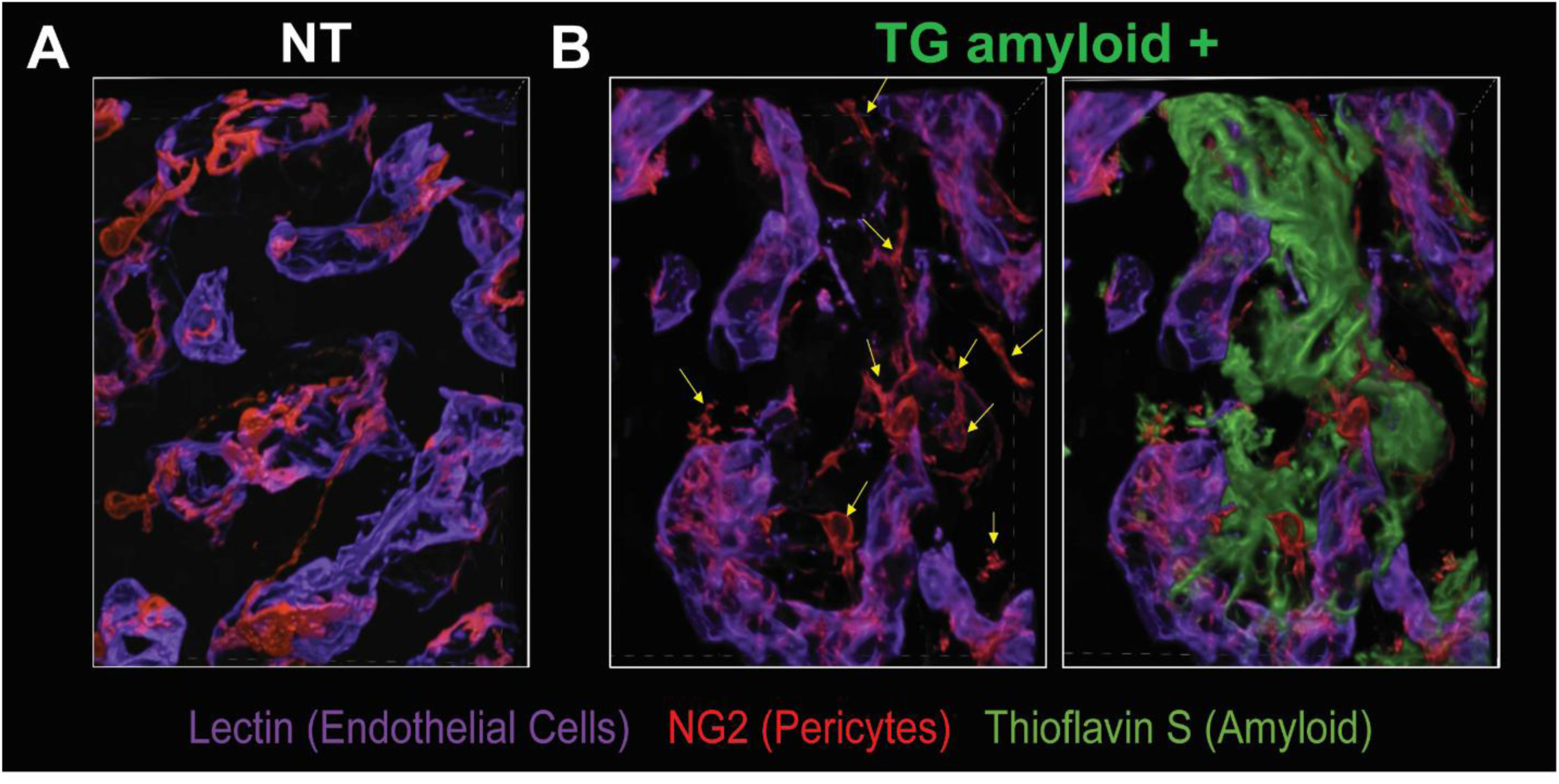
Representative volume projections of non-transgenic (NT) and hIAPP transgenic (TG) islet capillaries. A and B) 30um tissue sections were stained with lectin (purple), NG-2 (red), and Thioflavin S (green) and imaged with Nyquist sampling to generate 3D volume projections. Note that matched panels in A and B are the same visual field, but the green and grey channels (ThioflavinS, islet backscatter) were omitted for visualization purposes. Yellow arrows denote areas where NG2 staining is disconnected from lectin+ capillaries.

We next investigated the impact of hIAPP aggregation on microvascular morphology and function. In agreement with previous observations [13], we observed a decreased lectin-positive area in TG mice, with amyloid-positive islets exhibiting less uniformity and larger average diameter [**Fig 3A, 3B, 3D**]. TG microvessels in amyloid-negative islets were significantly narrower than NT vessels under basal conditions [**Fig 3C, 3D**], suggesting that hIAPP expression, even in the absence of aggregation, impacts islet vasculature. Additionally, islet pericyte area was significantly increased in TG animals, where pericytes and amyloid deposits were found in close contact with one another [**Fig 3E, 3F**]. Pericytes appeared to resist hIAPP-mediated cytotoxicity, and TG amyloid-positive islets had the highest pericyte density [**Fig 3F**].

**Figure 3:**
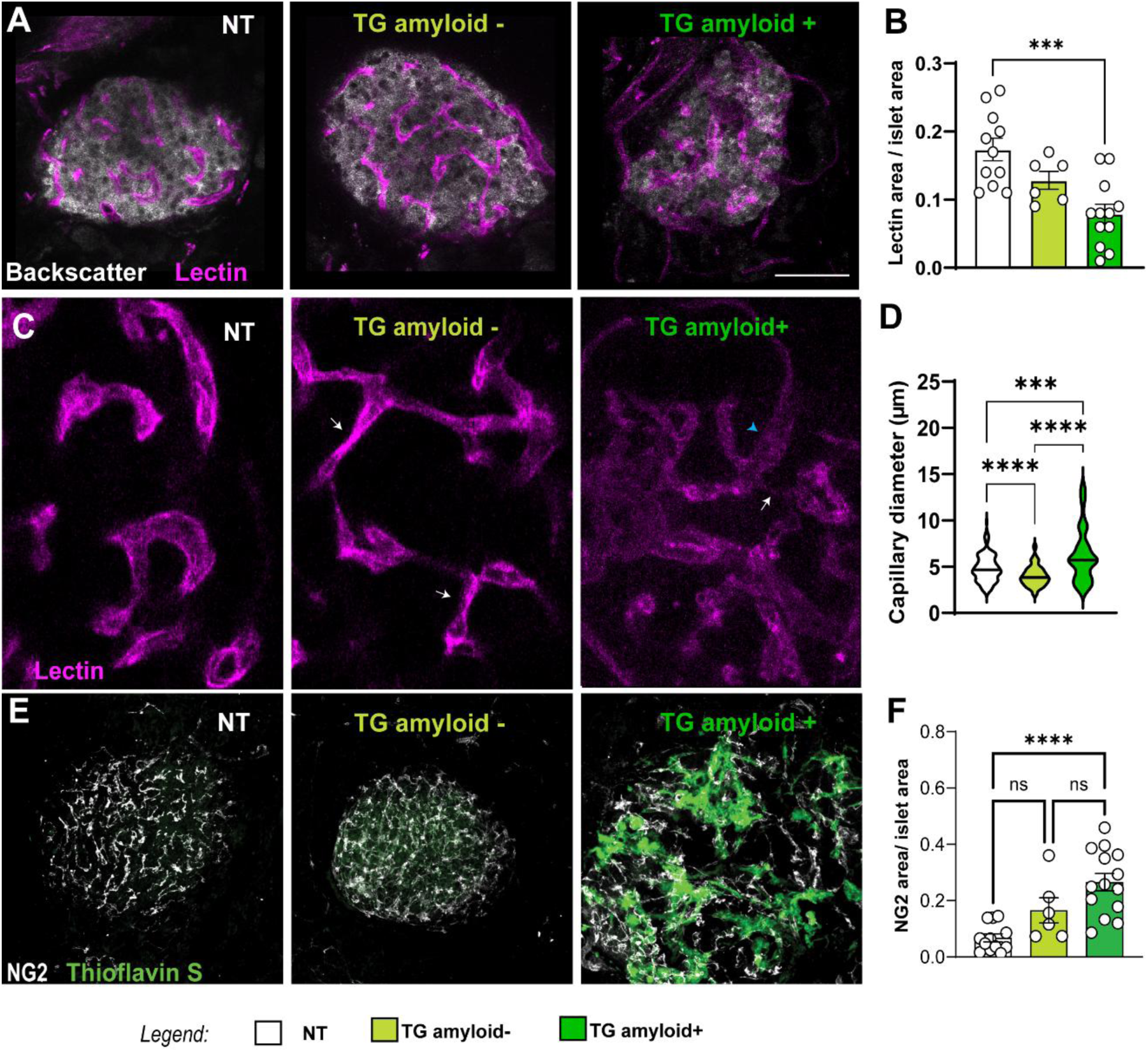
Endothelial cell and pericyte area in TG islets. A) Representative image of islet endothelial cells from NT and TG islets. B) Analysis of islet endothelial cell (lectin+) area across genotypes and amyloid status. C) Representative image of differences in capillary diameter where white arrows denote narrow vessels and blue arrowheads indicate enlarged vessels. D) Analysis of capillary diameter across genotype and amyloid status. E) Representative image of NG2+ pericytes from NT and TG islets. F) Analysis of islet NG2+ area across genotype and amyloid status. Vessel measurements were analyzed by Brown-Forsythe and Welch’s one-way ANOVA with Tukey-corrected multiple comparisons. Statistical significance is denoted by asterisks (* p<0.05, ** p<0.01, *** p<0.001, and **** p<0.0001).

Having characterized the structural differences between islet microvessels in the presence and absence of hIAPP aggregation, we next evaluated capillary function and pericyte Ca^2+^ dynamics in living pancreas slices [**Suppl Fig 5**]. We exposed *in situ* islet microvessels to vasomodulatory stimuli, including endothelin-1 (ET-1), bosentan (a dual ET-1 receptor antagonist), angiotensin II (AngII), epinephrine, and 16 mmol/l glucose. TG amyloid-positive capillaries showed no vasomotor responses to any stimulus applied, while NT and TG amyloid-negative capillaries only constricted in response to ET-1 or AngII, respectively [**Fig 4A; Suppl Fig 6**]. Despite their impaired vasomotion, we noted that TG amyloid-positive pericytes, as well as NT and TG amyloid-negative pericytes, mounted Ca^2+^responses to each vasoactive stimulus [**Fig 4B, 4C**]. TG amyloid-positive pericytes had the greatest Ca^2+^ response to both exogenous ET-1 and its receptor antagonist, bosentan, as determined by the AUC of fluorescence traces reflecting changes in intracellular Ca2+ levels elicited by each stimulus [**Fig 4B, 4C**]. In contrast, while Ca^2+^ responses to ET-1 or to bosentan were not significantly different from NT in TG amyloid-negative pericytes, these cells mounted a potent Ca^2+^ response to AngII [**Fig 4B, 4C**]. In summary, our data indicate that, in the presence of hIAPP aggregates, islet pericytes are dysfunctional and capillaries are unresponsive, potentially compromising blood flow regulation *in vivo*.

**Figure 4:**
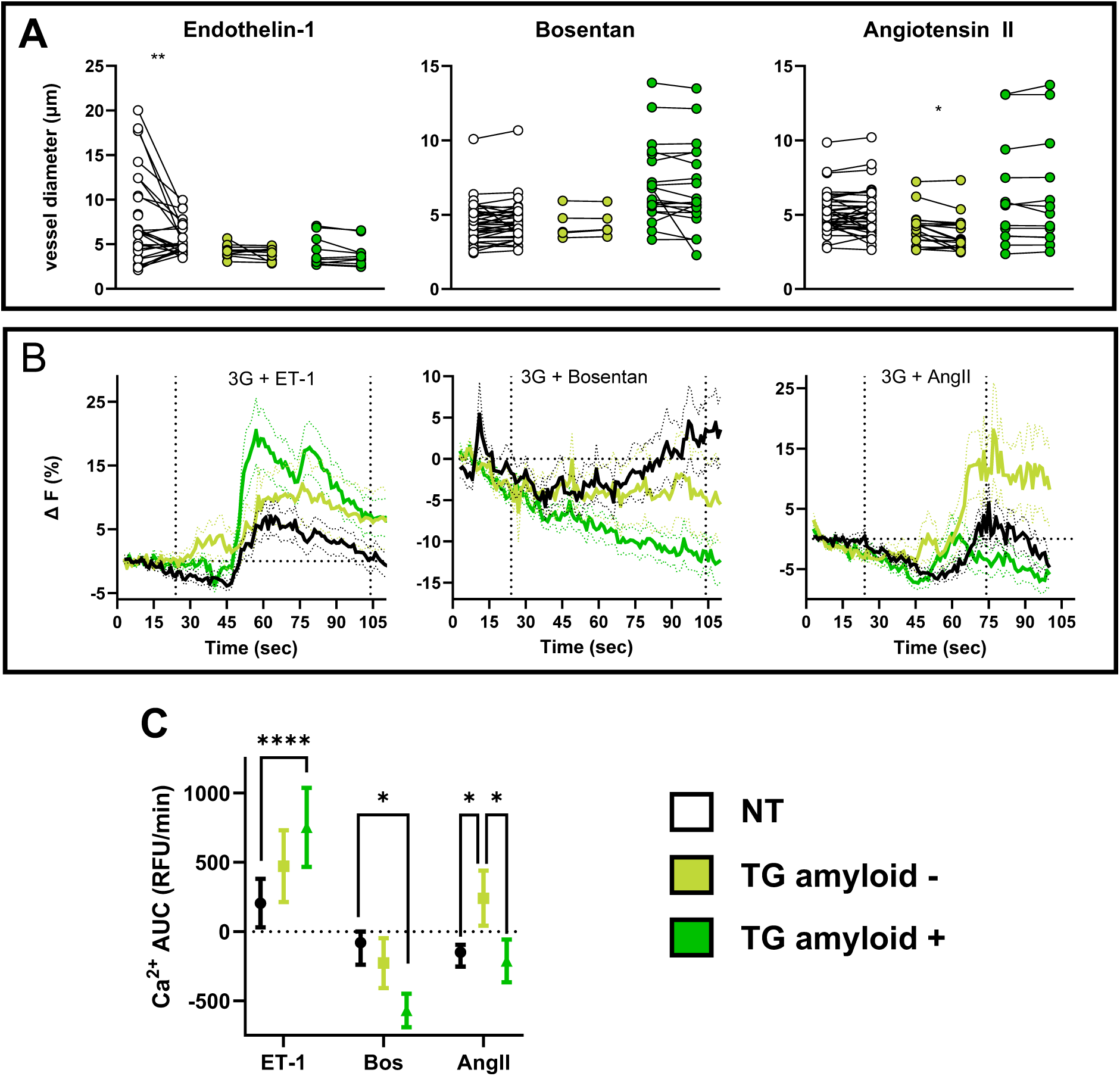
Live slice Ca2+ recordings from islet pericytes. A) Islet capillary diameter before and after Endothelin-1 (ET-1), bosentan (Bos), or Angiotensin-II (AngII) treatment. B) Change in fluorescence over time. C) Ca^2+^ AUC for the duration of each vasomodulatory stimulus. Panel A uses a two-way mixed-effects ANOVA with Šídák’s multiple comparisons. Panel C uses Brown-Forsythe and Welch’s one-way ANOVA with Tukey-corrected multiple comparisons. Statistical significance is denoted by asterisks (* p<0.05, ** p<0.01, *** p<0.001, and ****p<0.0001).

## DISCUSSION

Previous work has revealed decreased EC density and increased pericyte density adjacent to hIAPP deposits in TG mouse islets [13]. In the present study, we investigated transcriptional changes in remaining ECs and functional abnormalities in the expanded islet pericyte population. hIAPP-treated MS-1 cells and T2D human CD31+ islet ECs shared common transcriptional changes associated with impaired cell-cell adhesion and extracellular matrix formation, while functional recordings of live pancreatic slices revealed impaired vasomotion and altered pericyte Ca^2+^ dynamics. These data suggest that amyloid deposition not only compromises EC health and survival but also fundamentally alters islet microvascular morphology, stability, and function, potentially disrupting islet perfusion and exacerbating endocrine impairment in T2D.

### Endothelial cell transcriptional remodelling

Using GSEA and leading-edge gene analysis, we observed downregulation of a substantial number of GO pathways associated with the cytoskeleton and cell structure, ECM deposition and tissue remodelling, and cell-cell adhesion in both hIAPP-treated MS-1 cells and human CD31+ cells from T2D donors. These transcriptional programs suggest that vascular destabilization occurs early in the islet EC response to hIAPP and is present in T2D. Leading-edge gene analysis identified 35 common transcripts, with the top 3 being *Tln1, Thbs1,* and *Plec*. Of these, *PLEC* was the most downregulated leading-edge gene in human tissue, while *Tln1* was the most downregulated in MS-1 cells.

Talin-1 (*TLN1*, *Tln1*) is a ubiquitously expressed cytoskeletal and focal adhesion protein. In ECs, it is implicated in vascular integrity and resistance to capillary leakage [32, 33]. Related to talin-1, *PLEC* (Plectin, *Plec*) is involved in EC-EC junctional stability, providing stability in response to mechanical and shear stressors [34, 35]. If talin-1 or plectin were significantly downregulated in islet capillaries, their absence would contribute to vascular fragmentation at the level of EC-EC coupling. *Tln1* overexpression is associated with increased cancer metastasis and resistance to anoikis, a form of programmed cell death triggered by loss of anchorage, while *Tln1* depletion is associated with loss of cell-cell contact and impaired angiogenesis [32, 36, 37]. We observed decreased *Tln1* levels in hIAPP-treated MS-1 cells, which were Congo Red-attenuated. Feasibly, subtle reductions in *TLN1* could contribute to microvascular dysfunction, whereas substantial reductions would leave islet CD31+ cells more vulnerable to anoikis.

EC-derived thrombospondin-1 (*THBS1*, *Thbs1*) is both a membrane-anchored and secreted protein, with its function tightly regulated by extracellular peptidase and metalloprotease activity [38]. Membrane-anchored thrombospondin-1 regulates vascular remodelling by suppressing VEGF-mediated EC migration and inhibiting anoikis [39, 40]. Furthermore, extracellular thrombospondin-1 activates latent TGF-beta and contributes to downstream signalling processes in other cell types [41]. Islets lacking *Thbs1* demonstrate increased beta cell mass, but impaired beta cell function and reduced insulin biosynthesis. This phenotype can be rescued with latent TGF-beta activation [42], although it is worth noting that vascular stability and pericyte function were not studied in this model. We observed a decrease in MS-1 *Thbs1* mRNA after hIAPP exposure, along with a trend toward THBS1 downregulation in human CD31+ cells, warranting further investigation with a larger sample size. Future studies should consider the relationship between hIAPP aggregation and thrombospondin-1, focusing on protein levels and proximity to relevant signalling partners within the islet ECM. Should thrombospondin-1 depletion be a hallmark of hIAPP-mediated vascular destabilization and endocrine dysfunction, strategies to restore its downstream signalling programs could preserve islet homeostasis.

### Microvascular function and morphology

In this study, we confirmed decreased capillary area in amyloid-positive islets, along with abnormal capillary morphology. Altered capillary structure may indicate microvascular fragmentation, as previously suggested in human T2D islet samples [15], a phenomenon predicted to impair islet perfusion. We also documented increased NG2+ pericyte area, a phenomenon observed in other mouse models of islet dysfunction [13, 31]. Despite increased NG2+ density, pericytes in amyloid-positive islets were detached from their underlying capillaries and unable to mount proper vasomotor responses. Our study is the first to directly examine pericyte function *in situ* in hIAPP transgenic animals and to assess the relative impact of amyloid deposition on islet pericyte physiology.

Islets from TG islets had striking differences in their basal capillary diameter, where TG amyloid-negative capillaries were substantially narrower than amyloid-positive or NT vessels. In contrast, TG amyloid-positive capillary diameters were heterogeneous, with an average diameter much larger than that of NT vessels. We attribute these differences in vascular tone to changes in the function and attachment of neighbouring pericytes. Indeed, physiological pericyte detachment in angiogenesis is followed by vasodilation, while pericyte detachment in disease contributes to capillary hyperdilation [43]. As amyloid deposits form between ECs and pericytes, amyloid-adjacent capillaries likely become dilated from lost pericyte anchorage.

As cytosolic Ca^2+^ is required for pericyte contraction, elevated intracellular Ca^2+^ could promote capillary narrowing in TG amyloid-negative islets. Reactive oxygen species potently induce store-operated Ca^2+^ release from CNS pericytes [44], and hIAPP elicits oxidative stress in other cell types [13, 24]. Perhaps TG amyloid-negative islet pericytes experience oxidative stress from hIAPP oligomers and protofibrils – intermediate, cytotoxic, species of amyloid that form before dense aggregates [45]. In healthy capillaries, pericytes and ECs are gap-junctionally coupled, allowing the exchange of Ca^2+^ between cell types and buffering intracellular Ca^2+^ [46, 47]. Pericyte Ca^2+^ loading may be exaggerated by hIAPP toxicity on neighbouring gap junctionally-coupled ECs, such that both cell types may contribute to elevated pericyte cytosolic Ca^2+^.

We also observed exaggerated Ca^2+^ responsiveness in TG pericytes, evident by delta-F traces and Ca^2+^ AUC for each vasomodulatory stimulus. Despite being unable to mount a vasomodulatory response to either ET-1 or its antagonist, bosentan, TG amyloid-positive pericytes showed the largest increases and decreases in Ca^2+^ AUC to each stimulus, respectively. Loss of pericyte attachment to ECs may impair this Ca^2+^ buffering between cell types and contribute to the exaggerated Ca^2+^ response observed in TG amyloid-positive islet pericytes. Loss of gap junction and tight junction coupling not only impairs signalling but creates a physical barrier between cell types. Basement membrane thickening in capillaries of the retina, brain, and skeletal muscles impairs vasoactivity, although it is generally attributed to fibrosis and excess collagen deposition [8, 10, 48]. In diabetic retinopathy, basement membrane thickening reduces vascular compliance and compromises pericytes’ ability to regulate capillary tone [8]. In amyloid deposition, inelastic material between ECs and pericytes may similarly reduce vasoactivity, despite pericytes’ contractile machinery and Ca^2+^ dynamics remaining largely intact.

### Open questions and future directions

We observed vascular destabilization and impaired capillary vasoactivity in islets positive for hIAPP aggregation. At present, it is unclear whether pericyte dissociation from ECs results from hIAPP-induced weakened cell-cell adhesion, physical displacement by amyloid aggregation, ECM abnormalities, or a programmed adaptive response to vascular injury. Pericyte detachment is a required physiological process preceding angiogenesis, whereby the EC secretome is altered to favour pericyte migration away from underlying capillaries [43, 49, 50]. hIAPP oligomers, fibrils, or aggregates may initiate a hypoxic or acute injury response, which directly stimulates pericyte migration away from ECs. Perhaps pericyte dissociation is intended as a transient and protective response so that hIAPP-affected vessels can regenerate, but amyloid deposits likely impair their re-recruitment. Defects in pericyte recruitment could be explained by downregulation of EC *Thbs1*, as previously discussed, but also from depletion of *Ece1* (endothelin-1 converting enzyme) or *Ltbp4* (latent TGF beta binding protein), later entries in our top 32 common leading-edge gene list. ET-1 and TGF beta are important mediators of vascular stability; therefore, downregulation of their binding partners may alter the EC secretome, impairing pericyte reattachment [9, 43, 51]. Future investigations could use a coculture system to monitor the effects of hIAPP aggregation on pericyte-EC association. If the addition of thrombospondin-1, ET-1, or latent TGF beta attenuates hIAPP-mediated destabilization in this system, then correcting the expression of their associated genes (*Thbs1, Ece1, Ltbp4*) may represent a novel therapeutic strategy for improving islet microvascular function.

Given the complex and chronic nature of T2D and the expected donor-to-donor heterogeneity, we conducted GSEA and leading-edge gene analysisto compare human T2D samples with the murine hIAPP model. GSEA is a more sensitive and holistic approach that leverages the entire transcriptomic signal instead of a subset of differentially expressed genes [52]. Since our initial RNA-seq analysis and subsequent qPCR on isolated human islet CD31+ cells were underpowered, acquiring more T2D and non-diabetic human islet samples for analysis or mining larger-scale sequencing datasets could be beneficial for ascertaining whether our list of downregulated leading-edge genes is differentially regulated in the context of T2D ECs.

Although our dataset characterizes the hIAPP- and T2D-associated transcriptional changes in ECs, it lacks a similar analysis in islet pericytes. At present, there are no commercially available islet pericyte cell lines. Future studies could employ immortalized brain pericyte cell lines, bead-purified primary islet pericytes, or spatial transcriptomic approaches to investigate how pericytes respond to hIAPP and T2D-associated stressors. Furthermore, only male TG mice were used in this study, primarily due to the low incidence of hIAPP deposition in female TG mice [25]. Nonetheless, complementary experiments should consider the impact of hIAPP deposition in female islets to confirm the translatability of our findings to the broader population. Overall, our study provides important insight into the microvascular consequences of islet amyloid deposition, but the mechanisms of capillary destabilization and functional impairment remain to be determined.

Islet microvascular dysfunction is a known but underinvestigated contributor to T2D pathophysiology. The present study demonstrates that hIAPP potentiates islet capillary destabilization by altering EC mRNA expression and impairing EC-pericyte attachment. Specifically, we observed decreased capillary area and increased pericyte area, along with changes in basal capillary tone and physical displacement of pericytes. While islet pericytes maintained approximately normal Ca^2+^ dynamics, the presence of hIAPP deposits impaired their ability to modulate capillary tone. Therapeutic strategies that prevent hIAPP aggregation may help preserve islet microvascular function, but an alternative approach to correct the disrupted islet capillary microenvironment must be considered.

## Acknowledgements

The authors have no conflicts of interest, relationships or activities to disclose.

This work was supported by the Department of Veterans Affairs, VA Puget Sound Health Care System (Seattle, WA) Merit Review I01-BX004063 (RLHM), Canada Excellence Research Chair Award CERC-22-0023 (RLHM) and the Seattle Institute for Biomedical and Clinical Research. JJC was supported by NIH T32 HL007028. The study sponsors/funders were not involved in the design of the study; the collection, analysis, and interpretation of data; writing the report; and did not impose any restrictions regarding the publication of the report.

We acknowledge Drs. Steven Kahn and Sakeneh Zraika at VA Puget Sound Health Care System for helpful discussions during compilation of this manuscript.

---

**Supplementary Movie 1:** Volume projection of islet vasculature in an NT islet. Two-channel 40x projection where purple is lectin and red is NG-2 marking endothelial cells and pericytes, respectively. Note that no amyloid was detected in NT islets.

**Supplementary Movie 2:** volume projection of islet vasculature in a TG amyloid-positive islet. Three-channel 40x projection where purple is lectin, red is NG-2, and green is thioflavinS marking endothelial cells, pericytes, and amyloid deposits, respectively.

**Supplementary Movie 3:** volume projection of islet vasculature in a TG amyloid-positive islet with no ThioS channel. Two-channel 40x projection where purple is lectin and red is NG-2, marking endothelial cells and pericytes, respectively. Note that this is the same visual field as Supplementary Movie 2, but the green channel (ThioflavinS) was omitted for visualization purposes.

Supplementary Videos 1-3

*(animated 3D projection of resonance scanner z-stacks)*

Supplementary datasets 1-3

*(DeSeq2, GSEA, LEGA spreadsheets)*

**Supplementary Figure 1:**
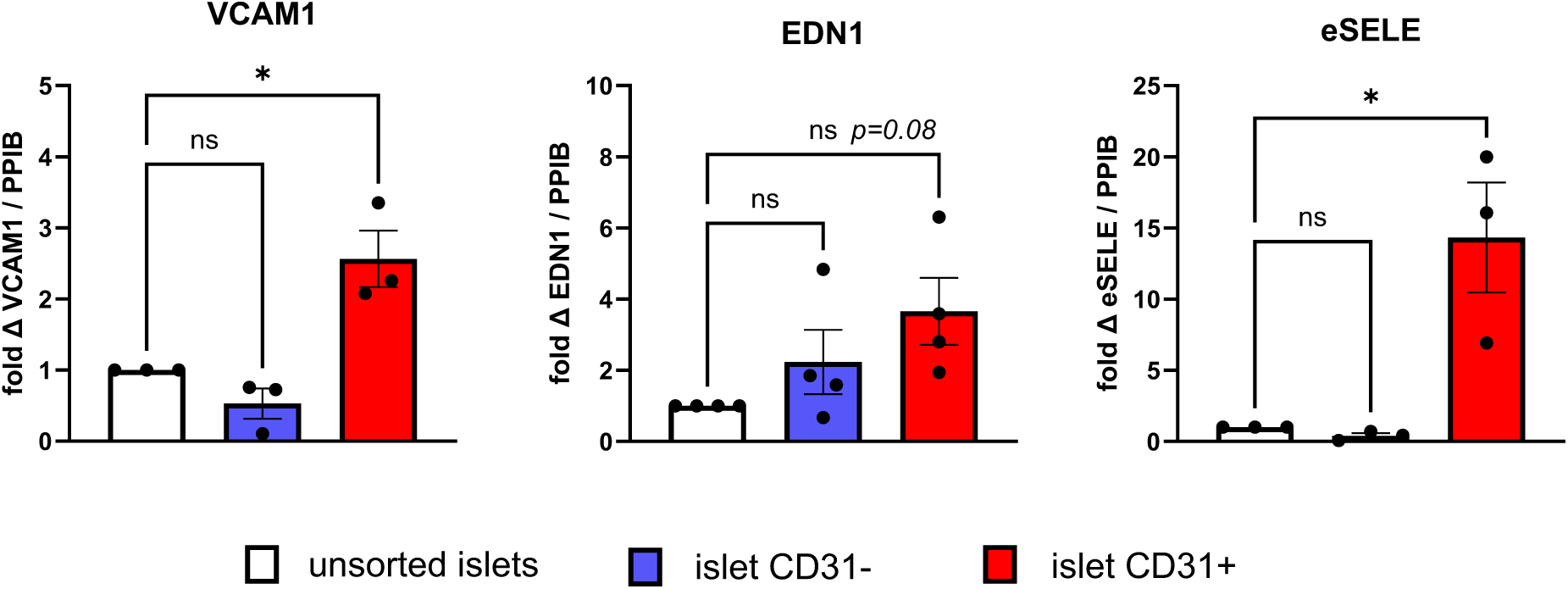
qPCR from donors HIE1, 7, 9, and 10, confirming endothelial cell enrichment after CD31+ magnetic bead sorting. All data are expressed as 2^-Δ ΔCt^ using the reference gene *PPIB* and normalized to unsorted islet mRNA. Statistics were determined by repeated measures one-way ANOVA, where statistical significance is denoted by asterisks (* p<0.05, ** p<0.01, *** p<0.001, and ****p<0.0001).

**Supplementary Figure 2:**
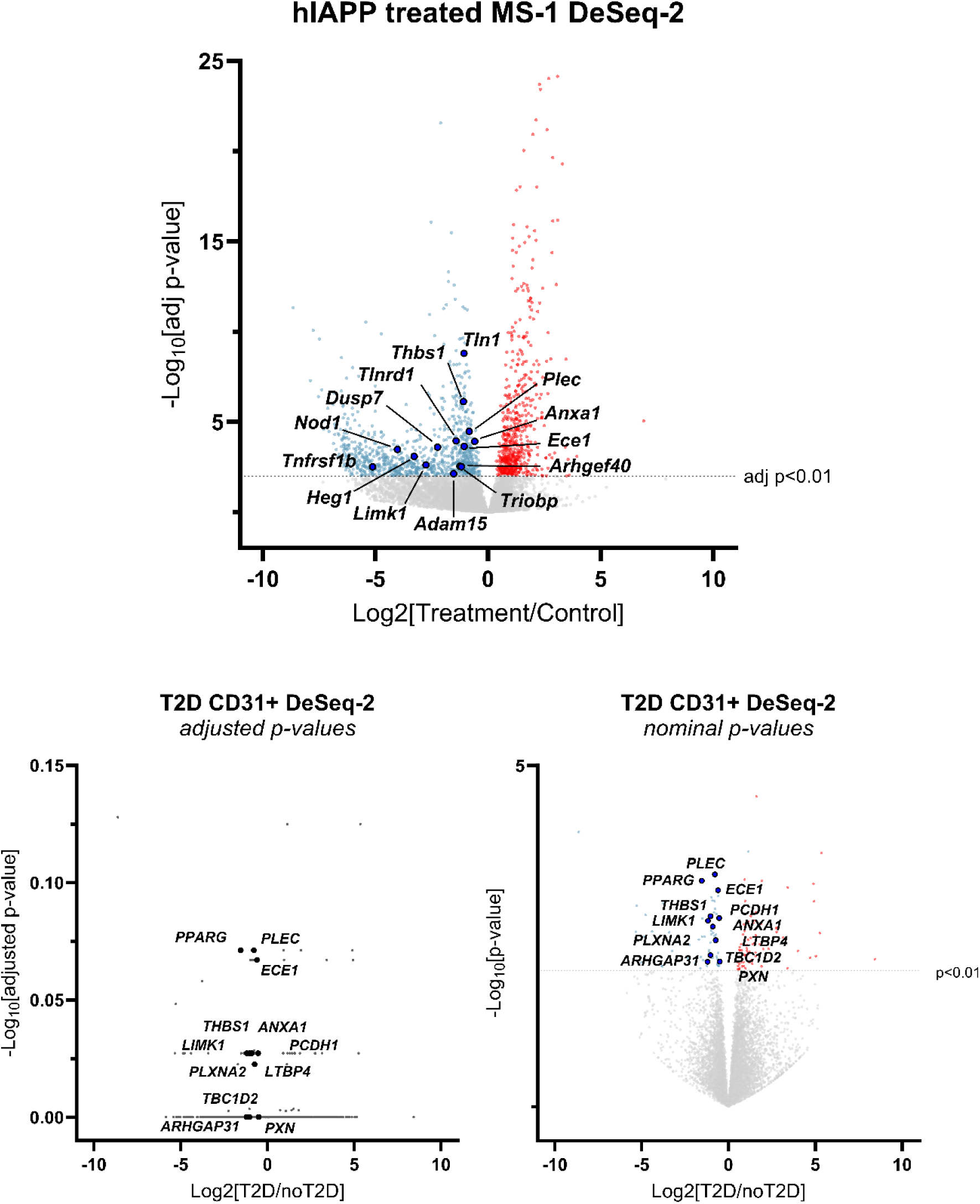
Volcano plots from MS-1 and human CD31+ cell DeSeq-2 analysis. A) differentially expressed genes in hIAPP-treated MS-1 cells. Labelled transcripts belong to the list of common leading-edge genes identified in Figure 1C. B and C) differentially expressed genes in bead-purified CD31+ cells from T2D islets plotted as adjusted (B) or nominal (C) p-values. Adjusted p-values were determined by Benjamini-Hochberg’s method.

**Supplementary Figure 3:**
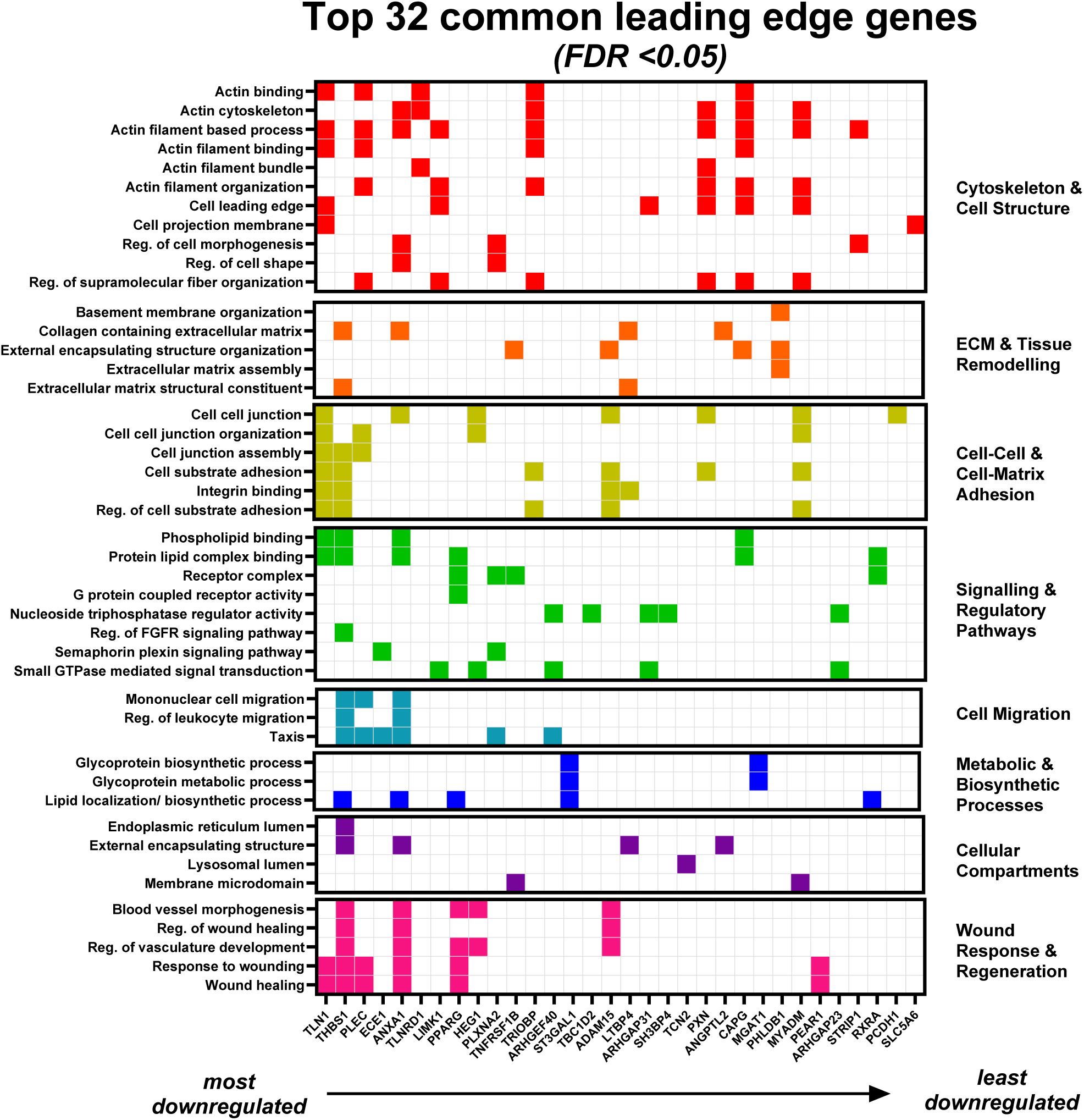
Expanded GSEA and Leading Edge Gene analysis. Binary heat map of common or equivalent GO processes identified in MS-1 and human CD31+ cells.

**Supplementary Figure 4:**
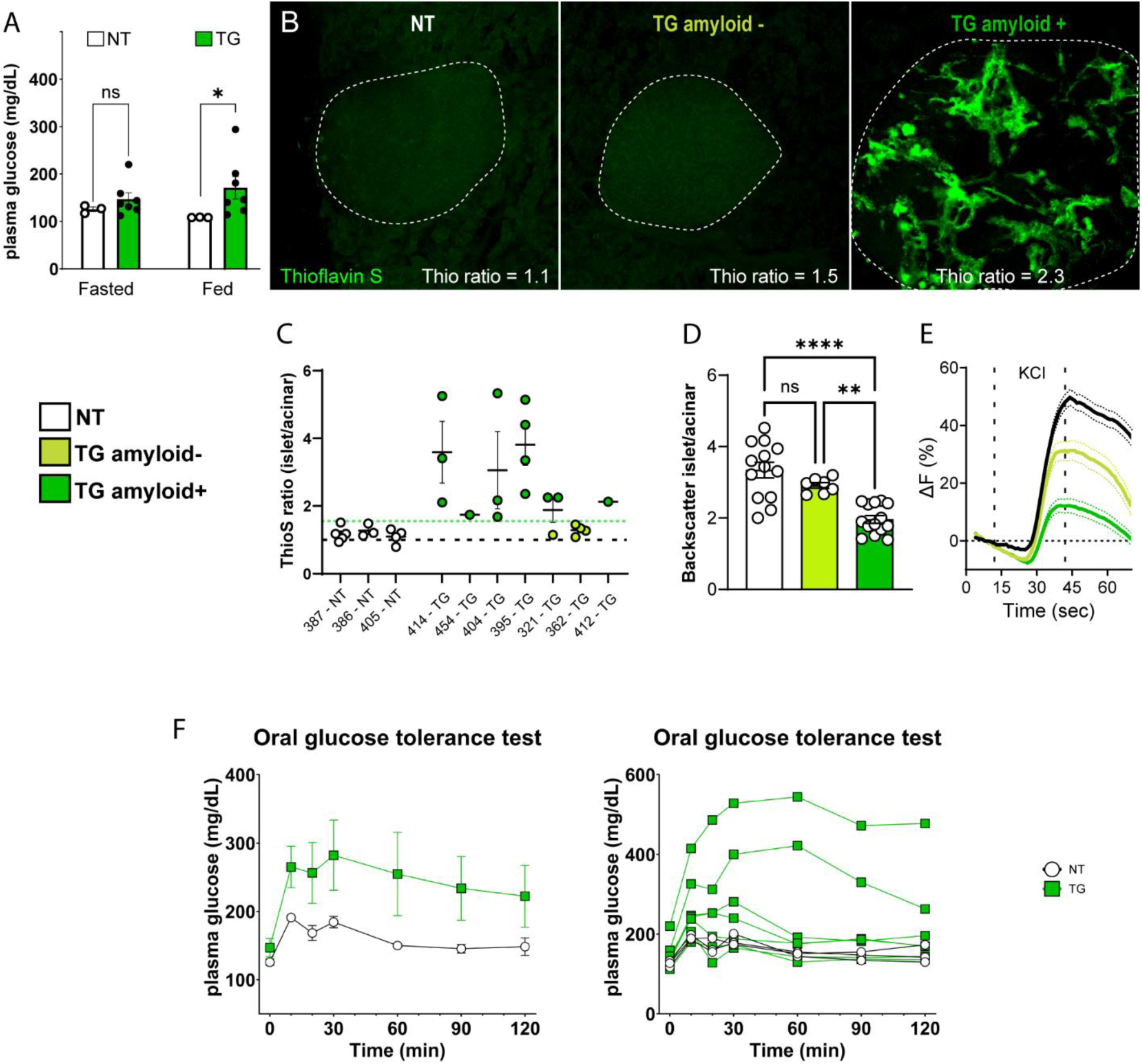
Amyloid and glucose tolerance phenotype of hIAPP transgenic mice. A) Fasted and fed plasma glucose measurements from NT and TG mice. B) Representative image of Thioflavin S staining, dashed outlines are islet perimeters defined by backscatter. C) ThioS ratio (islet ThioS fluorescence/acinar fluorescence), where the green dashed line is the cutoff for positive amyloid expression (3 standard deviations above the mean ThioS fluorescence of NT islets). D) Islet backscatter from NT and TG amyloid positive and negative islets, where amyloid positive islets have a ThioS ratio > 1.5, while amyloid negative islets have a ThioS ratio < 1.5. E) Change in total islet Ca^2+^ fluorescence (Fluo4) in response to KCl-mediated depolarization. F) Oral glucose tolerance test of N=3 NT and N=7 TG mice represented as mean ± SEM (left) or individual animals (right). Panel A uses Welch’s t-test. Panel D uses Brown-Forsythe and Welch’s one-way ANOVA with multiple comparisons. Statistical significance is denoted by asterisks (* p<0.05, ** p<0.01, *** p<0.001, and ****p<0.0001).

**Supplementary Figure 5:**
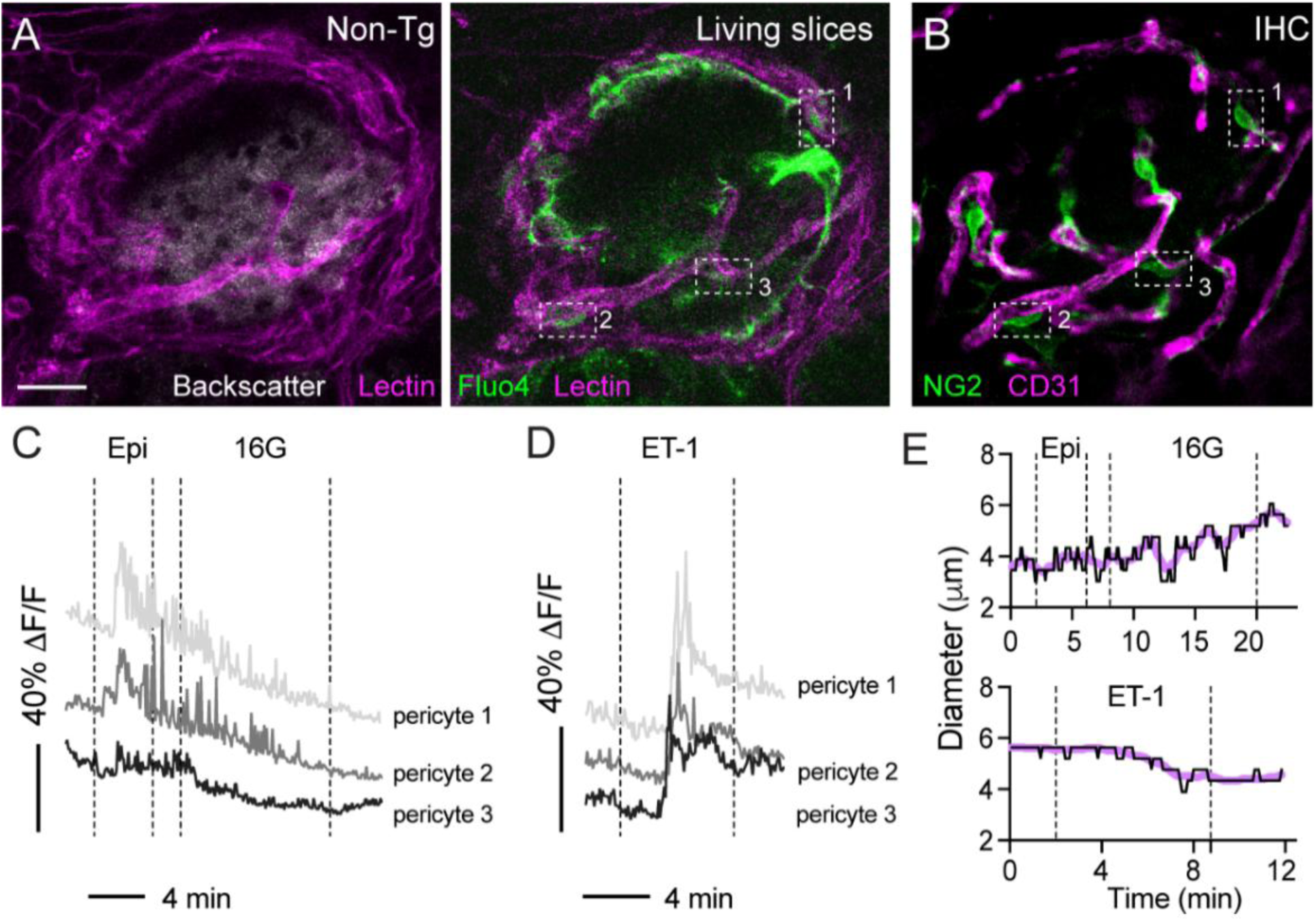
Sample image of live slice pericyte identification and recordings. A) Representative image of a live pancreatic slice from a lectin-infused NT mouse illustrating islet backscatter (white), lectin (purple) and fluo4 (green) fluorescence. Square callout boxes indicated presumed islet pericytes identified at the time of live-slice recording. B) Post hoc staining of the islet pictured in panel A, vasculature stained with CD31 (capillaries, purple) and NG2 (pericytes, green). Square callout boxes confirm the identity of islet pericytes. C and D) Sample fluorescence traces of 3 islet pericytes exposed to epinephrine (epi), 16mM glucose (16G), or endothelin-1 (ET-1). E) Sample diameter measurements of islet capillaries in response to epi, 16G, and ET-1.

**Supplementary Figure 6:**
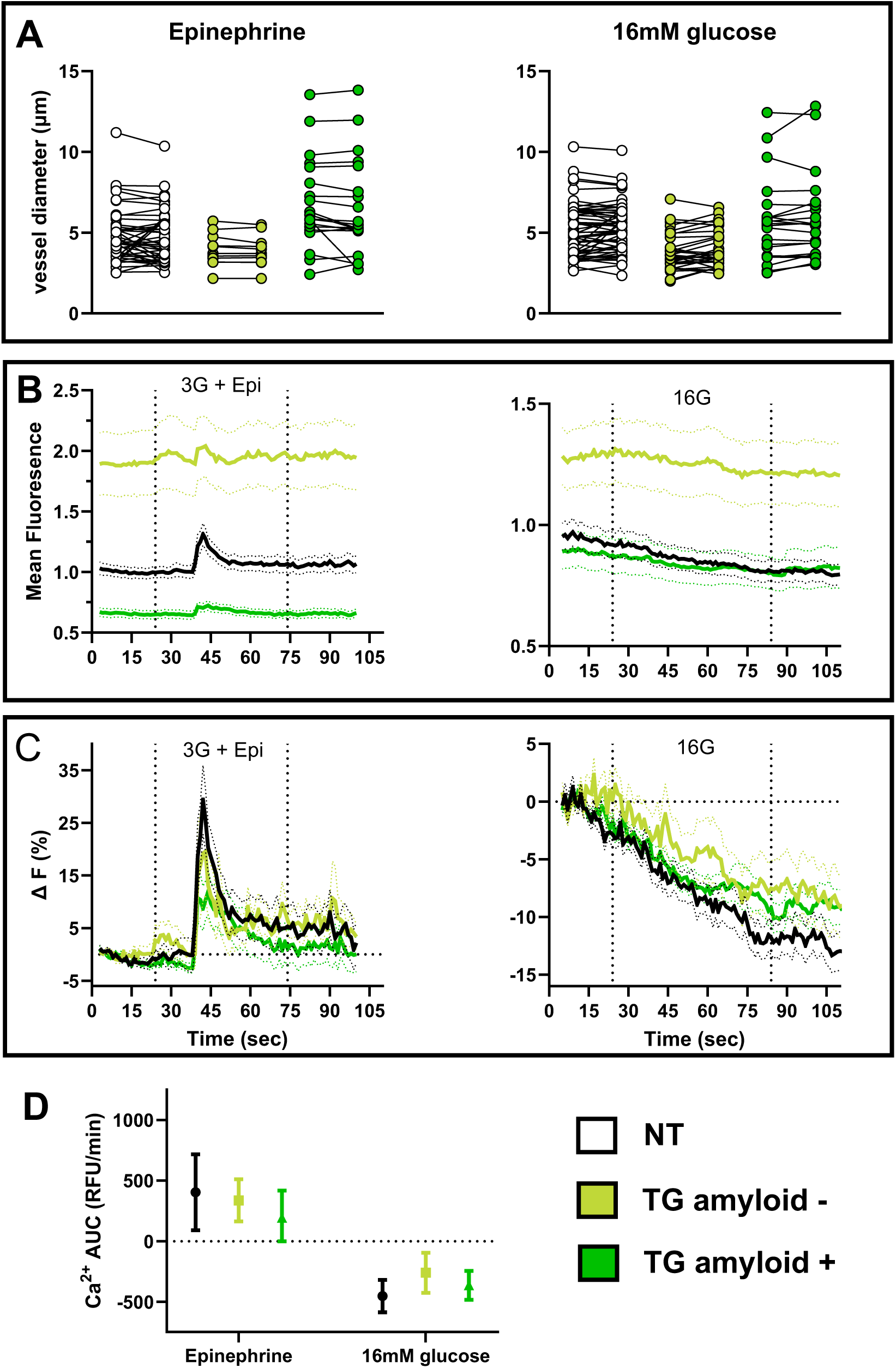
Live slice pericyte Ca^2+^ recordings with glucose and epinephrine. A) Islet capillary diameter before and after epinephrine of 16mM glucose exposure. B) Mean Ca^2+^ fluorescence normalized to islet ROI as defined by backscatter. C) Change in fluorescence over time. D) Ca^2+^ AUC for the duration of each vasomodulatory stimulus. Panel A uses mixed effects two-way ANOVA with Šídák’s multiple comparisons. Panel D use Brown Forscythe and Welch’s one-way ANOVA with multiple comparisons. All comparisons were non-significant.

**Supplementary Table 1:** Human islet donor characteristics.

| **Supplementary Table 1: Human islet donor characteristics** |  |  |  |  |  |  |  |
| --- | --- | --- | --- | --- | --- | --- | --- |
| Internal ID | RRID/ IIDPID | Diabetes status | HbA1c (%) | Age (years) | Sex | BMI (kg/m<sup>2</sup>) | Experiments |
| HIE1 | SAMN08769837 | T2 Diabetic | 6.6 | 48 | M | 43.7 | CD31+ qPCR |
| HIE3 | SAMN08769831 | Non-Diabetic | 5.5 | 51 | F | 20.1 | RNAseq |
| HIE4 | SAMN08769819 | T2 Diabetic | 6.6 | 45 | F | 30.5 | RNAseq |
| HIE7 | SAMN09228907 | Non-Diabetic | 5.8 | 62 | M | 36.1 | RNAseq, CD31+ qPCR, Islet qPCR |
| HIE8 | SAMN09370567 | Non-Diabetic | 5.2 | 32 | M | 28.5 | RNAseq, CD31+ qPCR |
| HIE9 | SAMN09432880 | Non-Diabetic | 5.4 | 51 | M | 35.7 | RNAseq, CD31+ qPCR |
| HIE10 | SAMN09644089 | Non-Diabetic |  | 27 | M | 32.9 | CD31+ qPCR |
| HIE11 | SAMN09844981 | T2 Diabetic | 7.6 | 56 | M | 29.4 | RNAseq, Islet qPCR |
| HIE12 | SAMN09862214 | Non-Diabetic | 5.3 | 31 | M | 31.8 | RNAseq |
| HIE13 | SAMN10372788 | T2 Diabetic | 9.9 | 50 | F | 35.4 | RNAseq, CD31+ qPCR, Islet qPCR |
| HIE14 | SAMN10832912 | T2 Diabetic | 8.0 | 54 | M | 33.7 | RNAseq |
| HIE15 | SAMN11157311 | T2 Diabetic | 7.3 | 34 | F | 31.7 | RNAseq, CD31+ qPCR |
| HIE16 | SAMN11606052 | T2 Diabetic | 10.1 | 58 | F | 34.8 | CD31+ qPCR, Islet qPCR |
| HIE27 | SAMN17831932 | T2 Diabetic | 7.5 | 60 | M | 41.3 | Islet qPCR |
| HIE28 | SAMN18021384 | Non-Diabetic | 5.8 | 56 | M | 24.2 | Islet qPCR |
| HIE29 | SAMN18092805 | Non-Diabetic | 5.1 | 56 | M | 21.6 | Islet qPCR |
| HIE30 | SAMN18200483 | Non-Diabetic | 5.4 | 48 | F | 47.6 | Islet qPCR |
| HIE34 | SAMN19920583 | Non-Diabetic | 5.4 | 54 | M | 24.5 | Islet qPCR |
| HIE46 | SAMN33293779 | T2 Diabetic | 11.5 | 49 | M | 68.5 | Islet qPCR |
| HIE55 | SAMN37680485 | T2 Diabetic | 6.7 | 63 | M | 33.9 | Islet qPCR |
| HIE56 | SAMN37737368 | Non-Diabetic | 5.8 | 50 | M | 38.6 | Islet qPCR |

**Supplementary Table 2:** TaqMan probes used for gene expression analysis.

|  |  |  |
| --- | --- | --- |
| **Gene Name** | **Species** | **TaqMan Probe ID** |
| 18S | Human | Hs99999901_s1 |
| Ppib (Cyclophilin) | Mouse | Mm00478295_m1 |
| Ppib | Human | Hs00168719_m1 |
| Thbs1 (Thrombospondin 1) | Mouse | Mm0049032_g1 |
| Thbs1 | Human | Hs00962908_m1 |
| Tln1 (Talin 1) | Mouse | Mm00456997_m1 |
| Tln1 | Human | Hs00196775_m1 |

